# Molecular basis for cysteine oxidation by Plant Cysteine Oxidases from *Arabidopsis thaliana*

**DOI:** 10.1101/2020.09.15.298976

**Authors:** Qiong Guo, Zhenzhen Chen, Gao Wu, Jie Wen, Shanhui Liao, Chao Xu

## Abstract

Plant Cysteine Oxidases (PCOs) play important roles in controlling the stability of Group VII ethylene response factors (ERF-VIIs) via N-Arg/degron pathway through catalyzing the oxidation of their N-Cys for subsequent Arginyl-tRNA--protein transferase 1 (ATE1) mediated arginine installation. Here we presented structures of PCO2, PCO4, and PCO5 from *Arabidopsis thaliana* (*At*PCOs) and examined their *in vitro* activity by MS. On the basis of Tris-bound *At*PCO2, we modelled the Cys-bound *At*PCO2 structure and identified key residues involved in N-Cys oxidation. Alanine substitution of potential N-Cys interaction residues impaired the activity of *At*PCO5 remarkably. The structural research, complemented by mutagenesis and mass spectrometry experiments, not only uncovers the substrate recognition and catalytic mode by *At*PCOs, but also sheds light on the future design of potent inhibitors for plant cysteine oxidases.

## Introduction

In Animals and plants, different mechanisms are exploited to sense oxygen concentration and regulate gene expression in response to low oxygen stress (hypoxia)[1]. Animals utilizes hypoxia-inducible transcription factor (HIF) to increase the expression levels of growth factors [2], while Plant Cysteine Oxidases (PCOs) play a critical role in oxygen homeostasis by serving as oxygen-sensing proteins[3, 4]. PCOs were found to control the turnover of Group VII ethylene response factors (ERF-VIIs) starting with Met-Cys-(^1^MC^2^), via N-degron pathway[4-7]. Once the N-terminal Met (N-Met) of target protein is excised by Met amino peptidase[8], the exposed N-terminal Cys (N-Cys) is subject to oxidization by PCOs, which covert the Cys to Cys-sulfinic acid (CysO_2_)[7]. After the oxidation of N-Cys, Arginyl-tRNA--protein transferase 1 (ATE1) installs an arginine before N-CysO_2_ to promote the proteasomal degradation through the Arg/N-degron pathway[4, 7].

The activity of PCOs were greatly compromised in the hypoxic condition, which promotes the expression of hypoxia-response genes by increasing the *in vivo* stability of the ERF-VIIs it mediated[5, 9]. Five ERF-VII family members were identified in *Arabidopsis thaliana*, including *At*RAP2.2, *At*RAP2.3, *At*RAP2.12, *At*HRE1, and *At*HRE2[5, 10]. The turnovers of above response factors mediated by PCOs provide a new layer in understanding how gene expression during stress response is regulated in a oxygen-dependent manner.

PCOs from *Arabidopsis thaliana* (*At*PCOs) has been found to catalyze the N-Cys oxidization of *At*HRE1 (^2^CGGAVIS^8^), *At*HRE2 (^2^CGGAIIS^8^), and *At*RAP2.2 (^2^CGGAIIS^8^), *etc* [6, 10]. Despite their important biological functions in response to low-oxygen stress, how PCOs recognize the substrate and catalyze the subsequent N-Cys oxidation remain elusive because they display low sequence identity with the dioxygenases that have known structures. Here we solved several crystal structures of PCOs from *Arabidopsis thaliana* (*At*PCOs), including those of *At*PCO2, *At*PCO4, and *At*PCO5. Although the three PCOs belong to two subfamilies, they display conserved catalytic pocket, suggesting the conserved catalytic mechanism among PCOs. Specifically, the structure of Tris-bound *At*PCO2 allows us to model an N-Cys in the catalytic pocket, which reveals the N-Cys specific recognition by PCOs. The PCOs-mediated N-Cys oxidization was complemented by mass spectrometry (MS) experiments. Therefore, our structure research not only provides structural insights into the N-Cys catalytic mechanism by PCOs, but also sheds light on the future design of chemical inhibitors in interfering with the activities of PCOs.

## Results

### *At*PCO2, *At*PCO4, and *At*PCO5 exhibit activities towards peptides derived from ERF-VII family members

To study the activities of *At*PCOs *in vitro* systematically, we cloned, expressed and purified the full-length or core fragment of several *At*PCOs, including *At*PCO2_48-276_, *At*PCO4_1-241_, and *At*PCO5_1-242_ (**Fig. 1A**). Then we synthesized two peptides derived from *At*HRE1 and *At*RAP2.2, *At*HRE1^2-10^ and *At*RAP2.2^2-8^, Respectively, and examine the activities of *At*PCOs towards the two peptides by MS experiments.

**Figure 1.**
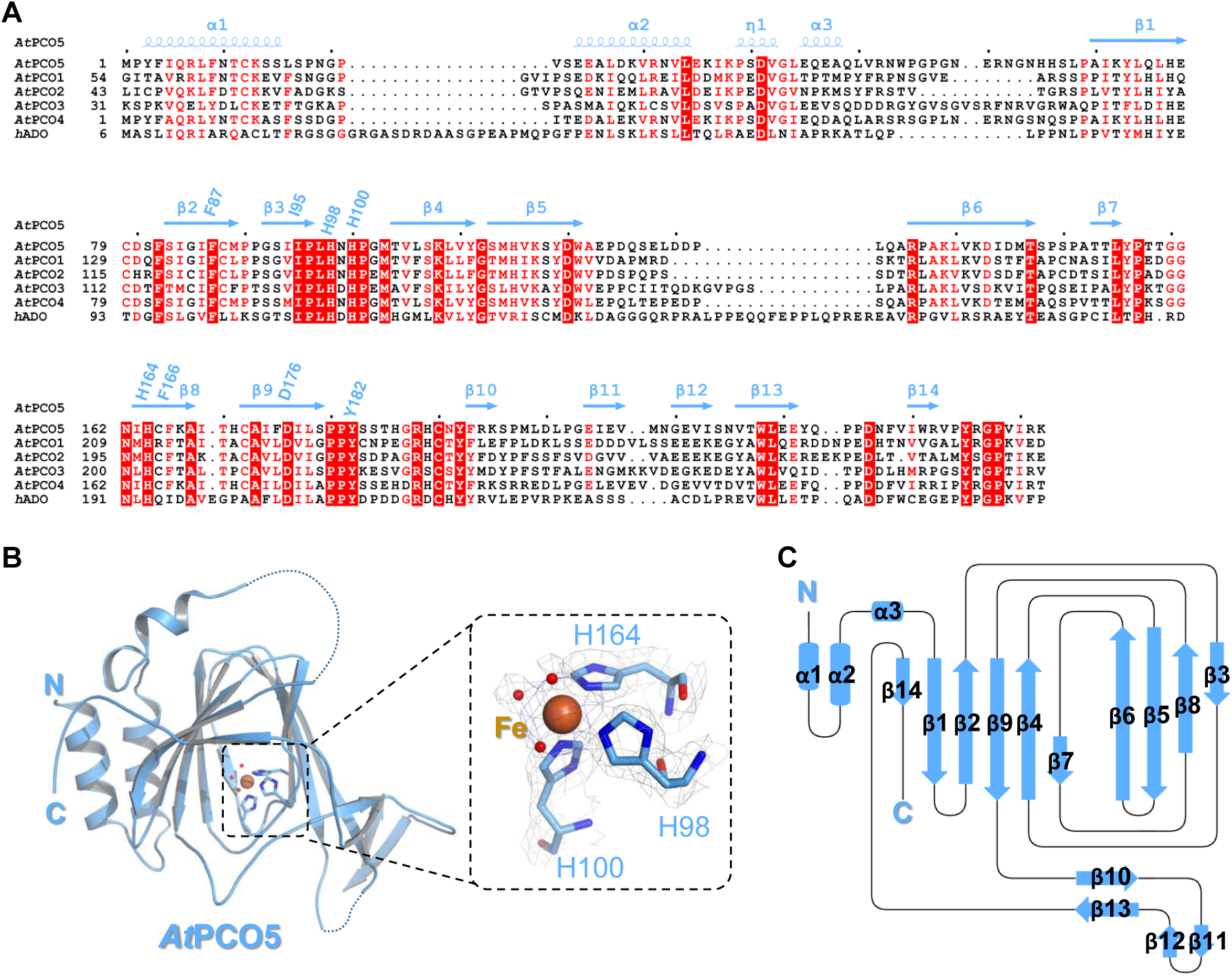
Structure of *At*PCO5. (A) Sequence alignment of *At*PCOs and human ADO (*h*ADO). Secondary structures of *At*PCO5 were labelled in blue at the top of aligned sequences. Three Fe^2+^ chelating histidines (His98, His100, and His164), and residues potentially involved in N-Cys recognition are labeled. (B) Overall structure of *At*PCO5_1-242_. *At*PCO5 is colored in blue, with three Fe^2+^ chelating histidines (His98, His100, and His164) shown in sticks. Fe^2+^ and three water molecules are shown in orange and red, respectively. (C) Topology diagram of *At*PCO5, with secondary structures colored in blue.

Consistent with previous reports about the control experiments[5, 11], after long incubation without enzyme, the peptides either dimerized via N-Cys (**Fig. 2A and 3A**) or formed thiazolidine (**Fig. 2B and 3B**). The dimerized peptide and formation of thiazolidine were confirmed by the peaks correspond to the ∼2-fold molecular weight and the +12 Da, respectively. In contrast, both peptides were efficiently catalyzed by *At*PCO2_48-276_, *At*PCO4_1-241_, and *At*PCO5_1-242_ in 30 minutes at 37°C, which generated a peak of +32 Da (**Fig. 2C-2E, and Fig. 3C-3E**)

**Figure 2.**
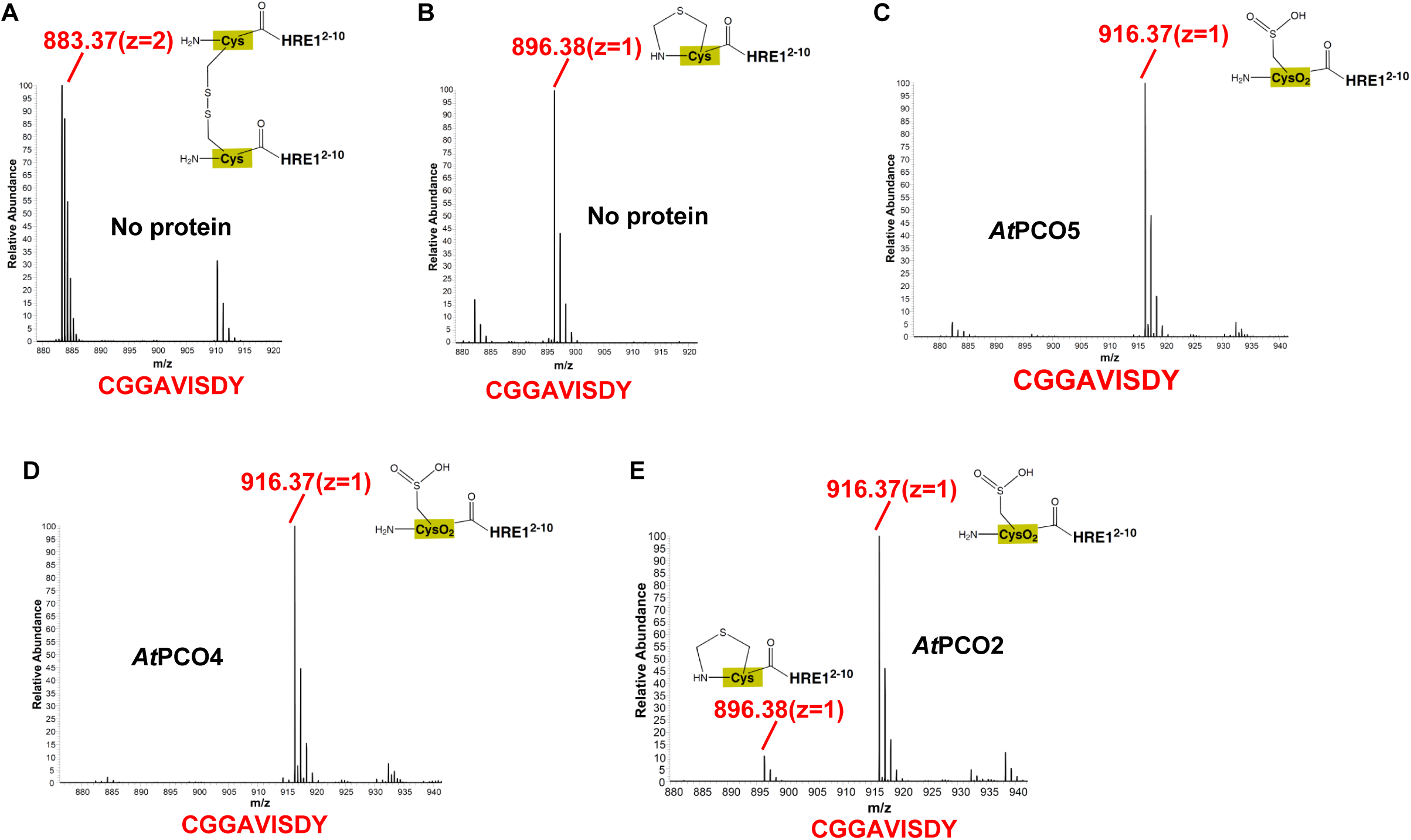
Mass spectra data of peptide *At*HRE1^2-10^ (CGGAVISDY) catalyzed by *At*PCOs. (A)-(B) Spectra of peptide without enzyme as the controls. Spectra the peptide CGGAIIS oxidized by (C) *At*PCO5_1-242_, (D) *At*PCO4_1-241_, and (E) *At*PCO2_48-276_.

**Figure 3.**
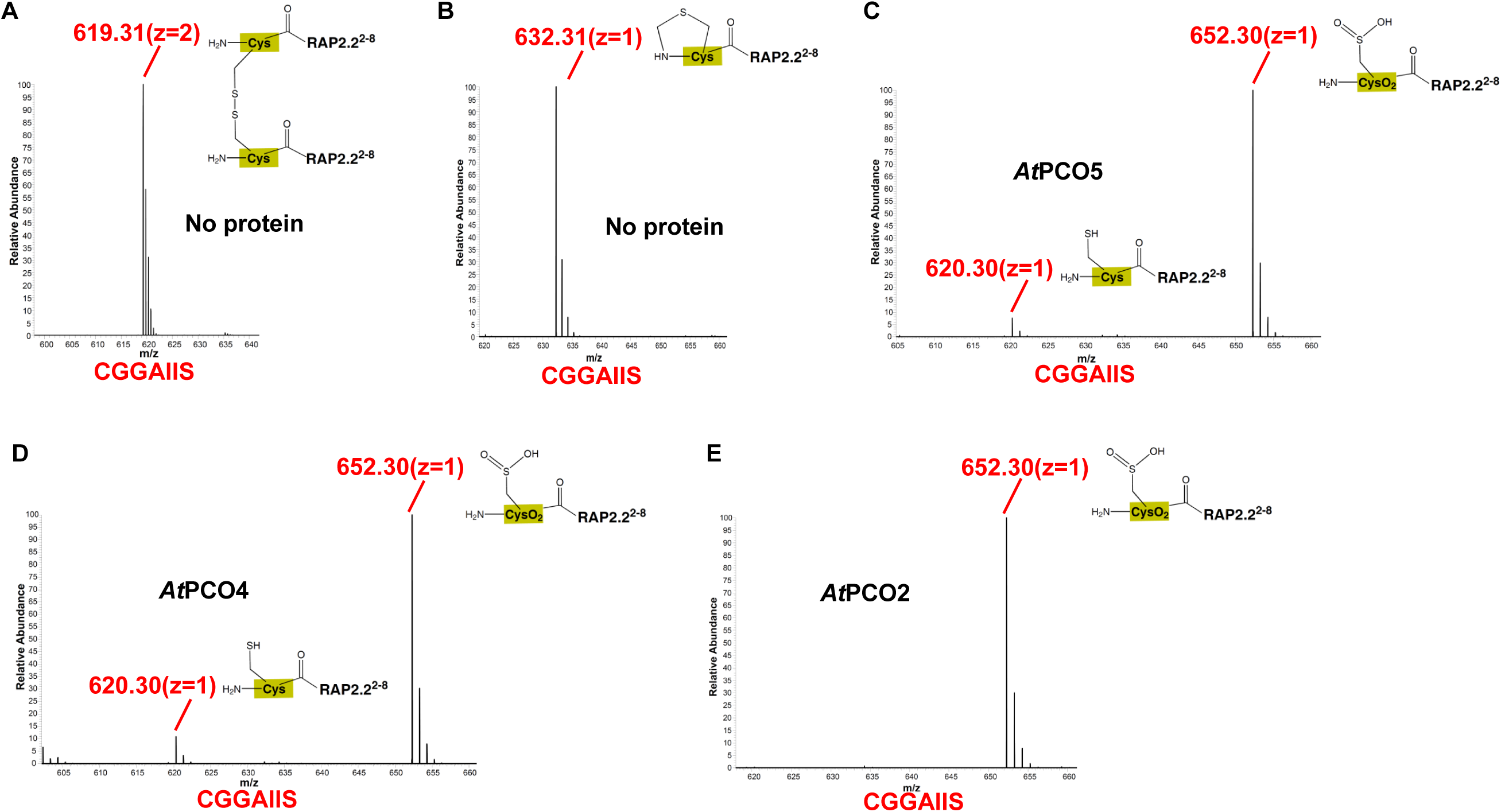
Mass spectra data of peptide AtRAP2.2^2-8^ (CGGAIIS) catalyzed by *At*PCOs. (A)-(B) Spectra of peptide without enzyme as the controls. Spectra the peptide CGGAIIS oxidized by (C) *At*PCO5_1-242_, (D) *At*PCO4_1-241_, and (E) *At*PCO2_48-276_.

### Crystal structure of *At*PCO5_1-242_, *At*PCO4_1-241_, and *At*PCO2_48-276_

To understand the substrate binding and catalytic mechanism of *At*PCOs, we solved the crystal structure of selenomethionine (SeMet) *At*PCO5_1-242_ at a resolution of 2.50 Å (**Table 1**). Most residues of *At*PCO5_1-242_ are visible except the residues at N-, C-terminus or some loop regions owning to internal flexibility (**Fig. 1B**). *At*PCO5 adopts a jelly roll-like fold with a chamber constituted by two β-sheets, β14-β1-β2-β9-β4-β7 and β6-β5-β8-β3, with the former β-sheet packed against α1-α3 (**Fig. 1B, 1C**). There is a long bended insertion between β9 and β14 consisting of β10-β13, in which β10 and β11 form anti-parallel strands with β13 and β12, respectively. Before crystallization, we added excess ferrous ion in the buffer and the density map of a ferrous ion is found in the catalytic center, which is chelated to His98, His100, His164, and three water molecules. His98 and His100 reside in the loop between β3 and β4, while His164 is localized in β8 (**Fig. 1**).

**Table 1.**
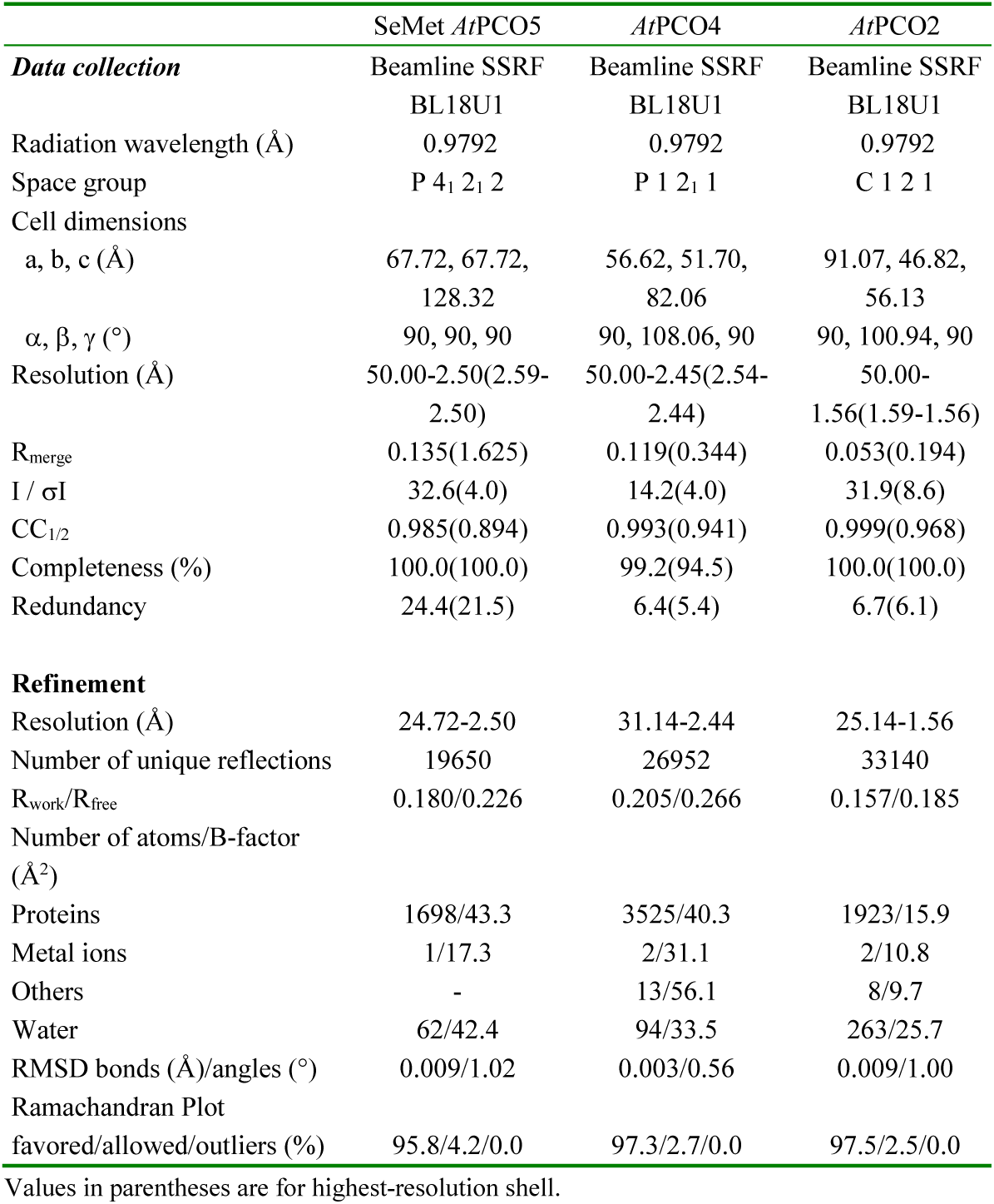
Data collection and refinement statistics.

|  | SeMet AtPCO5 | AtPCO4 | AtPCO2 |
| --- | --- | --- | --- |
| **Data collection** | Beamline SSRF | Beamline SSRF | Beamline SSRF |
|  | BL18U1 | BL18U1 | BL18U1 |
| Radiation wavelength (Å) | 0.9792 | 0.9792 | 0.9792 |
| Space group | P 4<sub>1</sub> 2<sub>1</sub> 2 | P 1 2<sub>1</sub> 1 | C 1 2 1 |
| Cell dimensions |  |  |  |
| a, b, c (Å) | 67.72, 67.72,<br>128.32 | 56.62, 51.70,<br>82.06 | 91.07, 46.82,<br>56.13 |
| $\alpha$, $\beta$, $\gamma$ (°) | 90, 90, 90 | 90, 108.06, 90 | 90, 100.94, 90 |
| Resolution (Å) | 50.00-2.50(2.59-2.50) | 50.00-2.45(2.54-2.44) | 50.00-1.56(1.59-1.56) |
| R<sub>merge</sub> | 0.135(1.625) | 0.119(0.344) | 0.053(0.194) |
| I / σI | 32.6(4.0) | 14.2(4.0) | 31.9(8.6) |
| CC<sub>1/2</sub> | 0.985(0.894) | 0.993(0.941) | 0.999(0.968) |
| Completeness (%) | 100.0(100.0) | 99.2(94.5) | 100.0(100.0) |
| Redundancy | 24.4(21.5) | 6.4(5.4) | 6.7(6.1) |
| **Refinement** |  |  |  |
| Resolution (Å) | 24.72-2.50 | 31.14-2.44 | 25.14-1.56 |
| Number of unique reflections | 19650 | 26952 | 33140 |
| R<sub>work</sub>/R<sub>free</sub> | 0.180/0.226 | 0.205/0.266 | 0.157/0.185 |
| Number of atoms/B-factor<br>(Å<sup>2</sup>) |  |  |  |
| Proteins | 1698/43.3 | 3525/40.3 | 1923/15.9 |
| Metal ions | 1/17.3 | 2/31.1 | 2/10.8 |
| Others | - | 13/56.1 | 8/9.7 |
| Water | 62/42.4 | 94/33.5 | 263/25.7 |
| RMSD bonds (Å)/angles (°) | 0.009/1.02 | 0.003/0.56 | 0.009/1.00 |
| Ramachandran Plot |  |  |  |
| favoured/allowed/outliers (%) | 95.8/4.2/0.0 | 97.3/2.7/0.0 | 97.5/2.5/0.0 |

Values in parentheses are for highest-resolution shell.

Next we solved structures of *At*PCO4_1-241_ and *At*PCO2_48-276_, at resolutions of 2.44- and 1.56-Å, respectively (**Fig. 4A, 4B and Table 1**). Both *At*PCO4 and *At*PCO2 adopt jelly roll-like fold (**Fig. 4A, 4B**). Superposition of three *At*PCO structures show that their overall architecture are very similar, as evidenced by the root-mean-square deviations (RMSDs) in a range of 0.57-0.81 Å (**Fig. 4C**). Especially the three histidines that are chelated to the ferrous ion are conserved in three structures, as an indicative of the conserved ferrous ion chelation mode in *At*PCOs (**Fig. 4A, 4B**). On the basis of sequence alignment, it is very likely that above (His)_3_ chelation mode also applies for their human homolog, 2-aminoethanethiol dioxygenase (ADO) (**Fig. 1A**).

**Figure 4.**
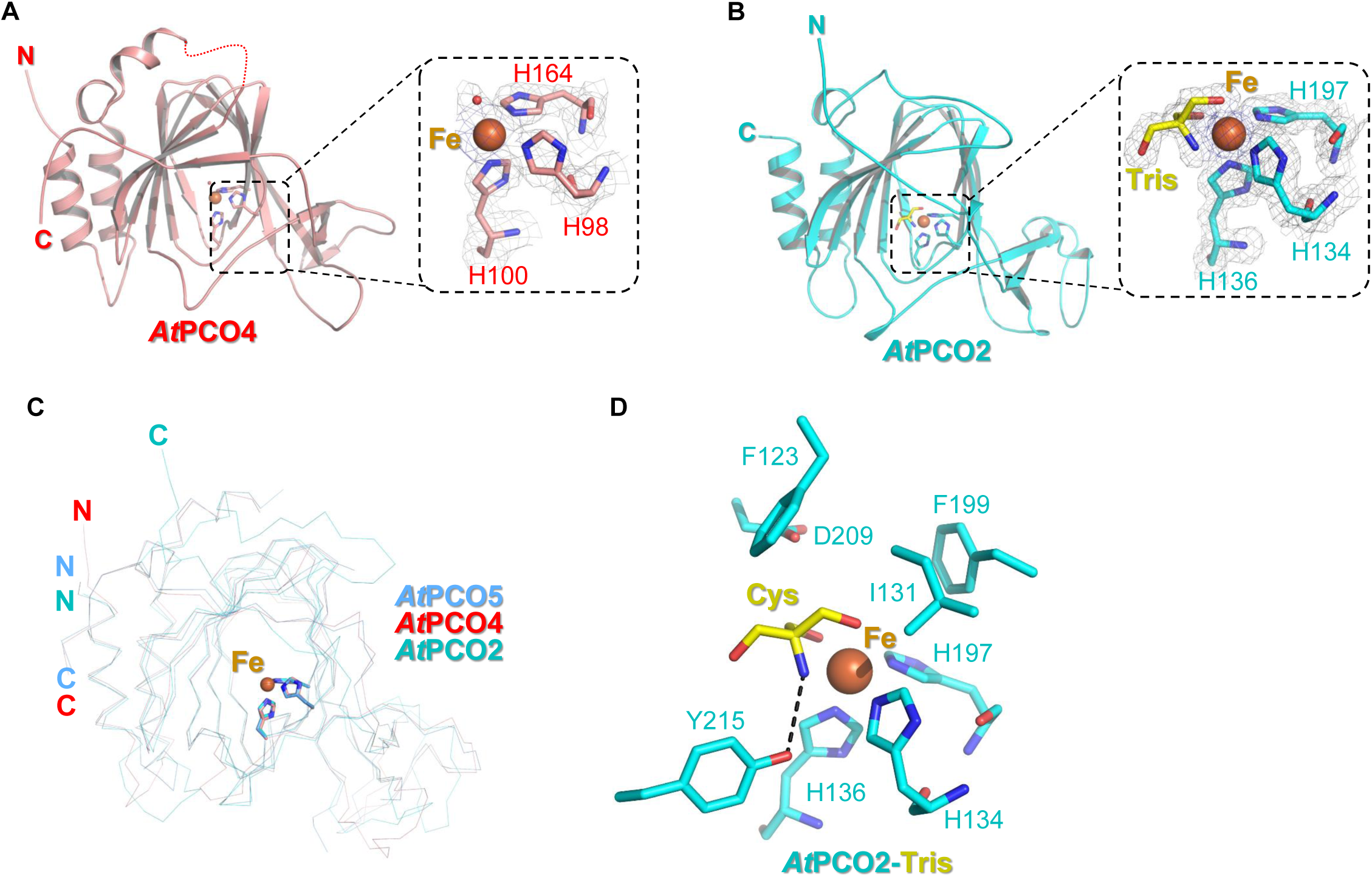
Structures of *At*PCO4_1-241_, and *At*PCO2_48-276_. (A) Overall structure of *At*PCO4_1-241_. The structure of *At*PCO4 is shown in red cartoon, with three Fe^2+^ chelating histidines (His98, His100, and His164) shown in sticks. Fe^2+^ and three water molecules are shown in orange and red, respectively. (B) Overall structure of Tris-bound *At*PCO2_48-276_. The structure of *At*PCO2 is colored in cyan cartoon, with three Fe^2+^ chelating histidines (His134, His136, and His197) shown in sticks. Fe^2+^ and Tris are shown in orange and yellow, respectively. (C) Superposition of the structures of *At*PCO2 (cyan), *At*PCO4 (red), and *At*PCO5 (blue). The structures are shown in ribbon, with Fe^2+^ chelation residues shown in sticks. (D) Detailed interactions between Tris and *At*PCO2. Tris is shown in yellow sticks. Fe^2+^ chelating residues and the Tris-binding residues are shown in cyan sticks. Intermolecular hydrogen bond is indicated by black dashes.

Intriguingly, in the structure of *At*PCO2, we found that a Tris molecule, mimic of the N-Cys, occupies the catalytic pocket. In the Tris-bound structure, besides the three histidines, the central ferrous ion is also chelated to the nitrogen and the two carboxyl groups of the Tris molecule to complete the six coordination (**Fig. 4D**). The nitrogen group of Tris is hydrogen bonded to the hydroxyl group of the *At*PCO2 Tyr215. Tris also makes van del Walls interactions with Phe123, Ile131, Phe199, and Asp209 of *At*PCO2 (**Fig. 4D**).

### Modeled structure of Cys-bound *At*PCO2

Given the similarity between the Tris and the apo cysteine, we modeled the Cys-bound structure of *At*PCO2 on the basis of the Tris-bound structure and compared its Cys recognition mode with that of recently solved Cys-bound human CDO (PDB ID: 6N42)[12]. Despite the similar architecture and Ferrous ion chelation mode, the jelly-roll fold of CDO deviates remarkably from that of *At*PCO2 (**Fig. 5A**). At the catalytic center, the modelled Cys is chelated to the Fe^2+^ via its main chain amino and side chain thiol groups, similar to that observed in the structure of Cys-bound CDO (**Fig. 5B, 5C**). Despite the similarity, their Cys recognition and catalytic modes are different in several aspects. Firstly, Cys93 and Tyr157 of CDO forms the cysteinyltyrosine bridge to increase the catalytic efficiency (**Fig. 5B**) [12, 13], whereas the crosslinked moiety is not found in any of *At*PCO structures (**Fig. 5C**). Secondly, the nitrogen group of Cys in the Cys-CDO complex is free, whereas the nitrogen group of Cys is hydrogen bonded to the side chain of Tyr215 in the modeled *At*PCO2 complex (**Fig. 5B, 5C**). Thirdly, in both structures, the Cys interaction residues in two structures are distinct (**Fig. 5B, 5C**). In the CDO structure, the Cys makes van der Waals interactions with the side chains of Leu75, Ser83, Val142, and His155 (**Fig. 5B**). In the modelled complex structure of *At*PCO2, the Cys makes van der Waals interactions with the side chains of Phe123, Ile131, and Phe199 of *At*PCO2 and its side chain makes one hydrogen bond with the side chain carboxyl group of Asp209, positioning the Cys in a favorable conformation for oxidation (**Fig. 5C**).

**Figure 5.**
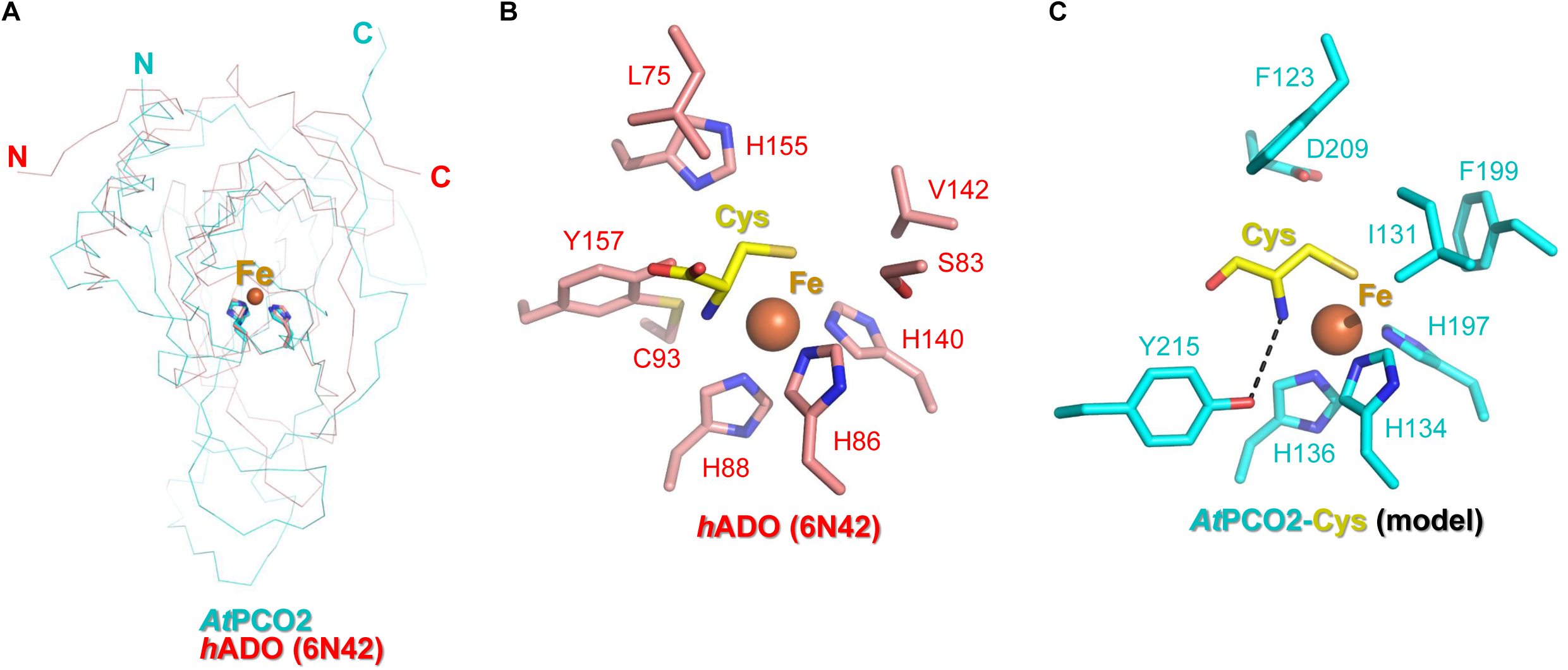
Comparison of the catalytic mechanism of *At*PCOs with that of human CDO. (A) Superposition of the structure of *h*CDO (red) with that of *At*PCO2 (cyan). The proteins are shown in ribbon, with Fe^2+^ chelating residues shown in sticks. (B) The catalytic center of Cys-bound *h*CDO, with the Cys and Cys binding residues shown sticks. (C) The catalytic center of Cys-bound *At*PCO2, with the Cys and Cys binding residues shown sticks.

The Cys interaction residues of *At*PCO2 are absolutely conserved in *At*PCO4 and *At*PCO5 (**Fig. 1A**), suggesting a common Cys recognition and catalytic mode shared by *At*PCOs. The counterparts of *At*PCO2 Asp209 and Tyr215 in *At*PCO5 are Asp176 and Tyr182, respectively (**Fig. 1A**). We then generated several single mutants for *At*PCO5_1-242_, including H164A, D176A, Y182F and C190A, and examined their activities towards the peptide RAP2.2^2-8^ (CGGAIIS). MS data show that while H164A abolished the activity by disrupting the ferrous chelation, both D176A and Y182F demonstrated compromised activities towards the *At*HRE1^2-10^ and *At*RAP2.2^2-8^ peptides than wild type *At*PCO5. In contrast, C190A exhibit comparable activity towards the substrate peptide as the wild type (**Supplementary Fig. S1**). Collectively, the mutagenesis experiments and MS data further confirmed the key roles of the Cys binding residues in cysteine oxidation.

## Discussion

Previous work on human CDO reveal that an cysteinyltyrosine bridge is formed to lower the oxidation potential of tyrosine for efficient catalysis[12, 13]. Whether cysteinyltyrosine bridge is also formed in human ADO and PCOs remains unknown because of the absence of structure evidence. By analyzing the structure of *At*PCOs, we proposed that although Cys190 of *At*PCO2 is spatially adjacent to Tyr215 and Tyr225 and the three residues are absolutely conserved in the structures of *At*PCO4 and *At*PCO5 (**Fig. 6**), it could not form the cysteinyltyrosine bridge with either of them. Firstly, the the cysteinyltyrosine bridge of CDO (Cys93-Tyr157) does not have a spatial counterpart in *At*PCO2. The Cys190 of *At*PCO2 is not the counterpart of CDO Cys92 in CDO **(Fig. 5B, 5C**). Secondly, the aromatic ring of *At*PCO2 Tyr225 is far from the thiol group of Cys223, implied that they are not likely to form crosslinked moiety. (**Fig. 6**). Thirdly, as for the Tyr215 of *At*PCO2, it is not likely to cross link with Cys223, either, because its aromatic ring of Tyr215 is parallel to the side chain plain of Cys223 and it would be energetically unfavorable for the Try215 to rotate its aromatic right at least ∼90 degree to form cysteinyltyrosine bridge. Also the main chain carbonyl group of Pro214 and the main chain amino group of Ser216 are hydrogen bonded to the side chain of Arg221 and the main chain carbonyl group of Arg221, respectively (**Fig. 6**), which does not allow the Tyr215 main chain to endure remarkable change. Above mentioned spatial pattern of the Cys223, Tyr215, and Tyr225 of *At*PCO2 are also conserved in *At*PCO4 and *At*PCO5 (**Fig. 6**), suggesting that *At*PCOs probably exploit a different mechanism rather than cysteinyltyrosine bridge to catalyze cysteine oxidation. We further mutating the Cys190 of *At*PCO4, the counterpart of the *At*PCO2 Cys223, to an Ala, and found that C190A does not impair the *At*PCO4 activity towards RAP2.2^2-8^ (**Supplementary Fig. S1**).

**Figure 6.**
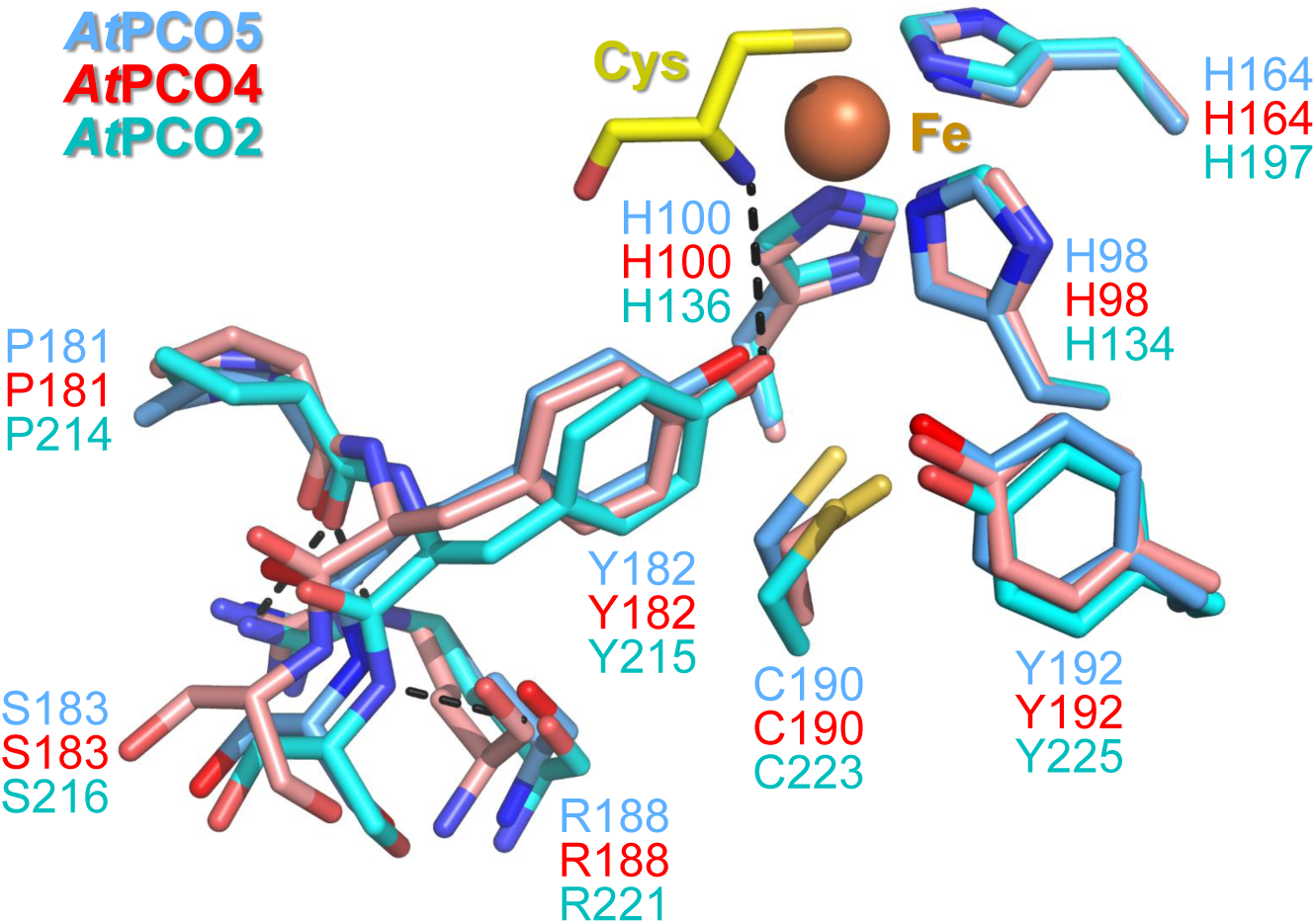
Spatial pattern of Cys223, Tyr215, and Tyr225 of *At*PCO2. Pro214 and Ser216 form main chain hydrogen bonds with Arg221. Fe^2+^ chelating residues, Arg221, Cys223, Pro214-Ser216, Tyr225, as well as their counterparts in *At*PCO4 and *At*PCO5, are shown in sticks.

During the preparation of the manuscript, the structures of *At*PCO4 and *At*PCO5 from *Arabidopsis thaliana* were reported[14]. In that study, the presented structures, as well as the conclusions drawn on the basis of the structures, are similar to ours. In near future, the substrate- or inhibitor-bound structures of *At*PCOs would further uncover the catalytic mechanism of N-Cys oxidation.

## Conclusion

Plant Cysteine Oxidases from Arabidopsis thaliana have been reported to play an important role in controlling the turnover of response factors via Arg/N-degron pathway. Our presented structures of *At*PCO2, *At*PCO4, and *At*PCO5 unraveled the architecture of PCOs, and the structure of ligand-bound *At*PCO2 revealed the potential substrate binding and oxidation residues in the catalytic pocket. Besides, sequence alignment indicates that human ADO might utilize a mechanism to catalyze the oxidation of N-Cys similar to those of *At*PCOs. Overall our research not only provides insight into the N-Cys oxidation mechanism by PCOs, but also sheds light on the design of inhibitors for PCOs.

## Acknowledgments

We thank the staff at beam line BL18U1 of the Shanghai Synchrotron Radiation Facility for providing technical support and assistance in data collection and analysis. This work was supported by the “Strategic Priority Research Program” of the Chinese Academy of Sciences (Grant No. XDB19000000) and National Natural Science Foundation of China Grants (31770806, 31500601). C. X. is also supported by the Major/Innovative Program of the Development Foundation of the Hefei Center for Physical Science and Technology (2018CXFX007) and the “Thousand Young Talent program”.

## Declarations of interest

The authors declare that they have no conflict of interest

## Author contributions

Q.G. and C.X. designed and conceived the study. Q.G., Z.C., S.L. purified, crystallized the protein, solved the structure, and analyzed the data. Q.G. and G.W. performed MS experiments. C.X. wrote the paper with the help from all authors. J.W., S.L., and C.X. supervised the experiments.

## Materials and Methods

### Cloning, mutation, protein expression and purification

Genes encoding full-length *At*PCO2, *At*PCO4, and *At*PCO5 were synthesized by Sangon Biotech (Shanghai). *At*PCO2_48-276_, *At*PCO4_1-241_, and *At*PCO5_1-242_ was amplified by polymerase chain reactions (PCR), and cloned into pET28-MHL (Genbank accession number: EF456735). The plasmids were then transformed into *E. coli* BL21 (DE3) and the recombinant proteins were overexpressed at 16°C for 20 h in the presence of 0.2 mM isopropyl b-D-1-thiogalactopyranoside (IPTG).

Cells were harvested at 3600 × g, 4 °C for 15 min, and then were resuspended using a buffer containing 20 mM Tris-HCl (pH 7.5), 400 mM NaCl (suspension buffer) and were lysed by sonication. Lysates were centrifuged at 18000 × g, 4 °C for 30 min and supernatants were collected. Recombinant proteins were purified with a fast flow Ni-NTA column (GE Healthcare) and eluted by 20 mM Tris-HCl (pH 7.5), 400 mM NaCl, 500 mM imidazole. Gel filtration and ion-exchange were employed for further purification. Gel filtration experiments were performed on a HiLoad™ 16/ 600 superdex™ 75 pg column (GE healthcare) with suspension buffer, the fractions containing target recombinant proteins were collected and dialyzed to ion exchange buffer A (20 mM Tris-HCl pH 7.5, 50 mM NaCl). Ion exchange experiments were performed on a Hitrap™ Q HP (1 mL) column (GE healthcare) with ion exchange buffer A and ion exchange buffer B (20 mM Tris-HCl pH 7.5, 1 M NaCl), fraction corresponding to target proteins were collected and concentrated to 20-40 mg/mL and stored at −80°C before further use. Seleno-Methionine (SeMet)-labeled *At*PCO5_1-242_ was purified in the same way, except that cells were cultured in M9 medium supplied with 50mg/L Seleno-Methionine. The mutants were constructed by conventional PCR using a MutanBEST kit (TaKaRa) and further verified by DNA sequencing. The mutants were expressed and purified in the same way as the wild type proteins.

### Crystallization, data collection and structure determination

Before crystallization, recombinant proteins were pre-incubated with Fe^2+^ at a molar ratio of 1:3 at 4 °C for 30 min. For crystallization of *At*PCOs, 1 μl protein was mixed with 1 μl crystallization buffer using the sitting drop vapor diffusion method at 18 °C. SeMet-labeled *At*PCO5_1-242_ was crystallized in a buffer containing 0.1 M MES monohydrate, pH 6.5, 12% (w/v) PEG20000. *At*PCO4_1-241_ was crystallized in a buffer containing 0.1 M Sodium citrate tribasic dehydrate, pH 5.0, 0.2 M Ammonium acetate, and 20% (w/v) PEG 3350. *At*PCO2_48-276_ was crystallized in a buffer containing 0.1 M Tris hydrochloride, pH 8.5, 0.2 M sodium acetate, and 30% w/v polyethylene glycol 4000. Before flash-freezing crystals in liquid nitrogen, all crystals were soaked in a cryo-protectant consisting of 90% reservoir solution plus 10% glycerol.

The diffraction data were collected on beam line BL18U1 at the Shanghai Synchrotron Facility (SSRF). Data sets were collected and processed using the HKL3000 program[15]. The initial model of SeMet *At*PCO5_1-242_ was solved by CRANK2[16], built manually by COOT[17] and refined by Phenix[18]. The structures of *At*PCO2_48-276_ and *At*PCO4_1-241_ were solved by molecular replacement by using the structure of SeMet AtPCO5_1-242_ as the search model. The structures were built manually by Coot[17] and were further refined by Phenix [18].

The atomic coordinates and structure factors of *At*PCO2, *At*PCO4, and *At*PCO5 have been deposited in the Protein Data Bank with PDB ID codes 7CHJ, 7CHI, and 7CXZ.

### Mass spectrometry experiments

Reversed-phase microcapillary/tandem mass spectrometry (LC/MS/MS) was performed using an Easy-nLC nanoflow HPLC (Proxeon Biosciences) with a self-packed 75 mm 15 cm C18 column connected to a QE-Plus (Thermo Scientific) in data-dependent acquisition and positive ion mode at 300 nL/min. Passing MS/MS spectra were manually inspected to ensure that all b-and y-fragment ions aligned with the assigned sequence and modification sites. A 25 μl reaction mixture contained 1 μM *At*PCO2/4/5 (final concentration) and 50 μM peptide (final concentration) in a buffer containing 20 mM Tris-HCl (pH 8.0), 20 mM NaCl, 20 μM FeSO _4_, 5 mM TCEP, and 1 mM Ascorbic acid. The reaction was incubated at 37°C for 30min at atmospheric oxygen before being quenched (at 70°C for 10–15 mins). Then, the tubes were centrifuged at 15000×g for 10 min. Supernatants were analyzed by LC-MS/MS and Proteomics Browser software, with the relative abundances of substrate and product reflecting the cysteine oxidation activities of proteins.

---

**Supplementary Fig. S1.**
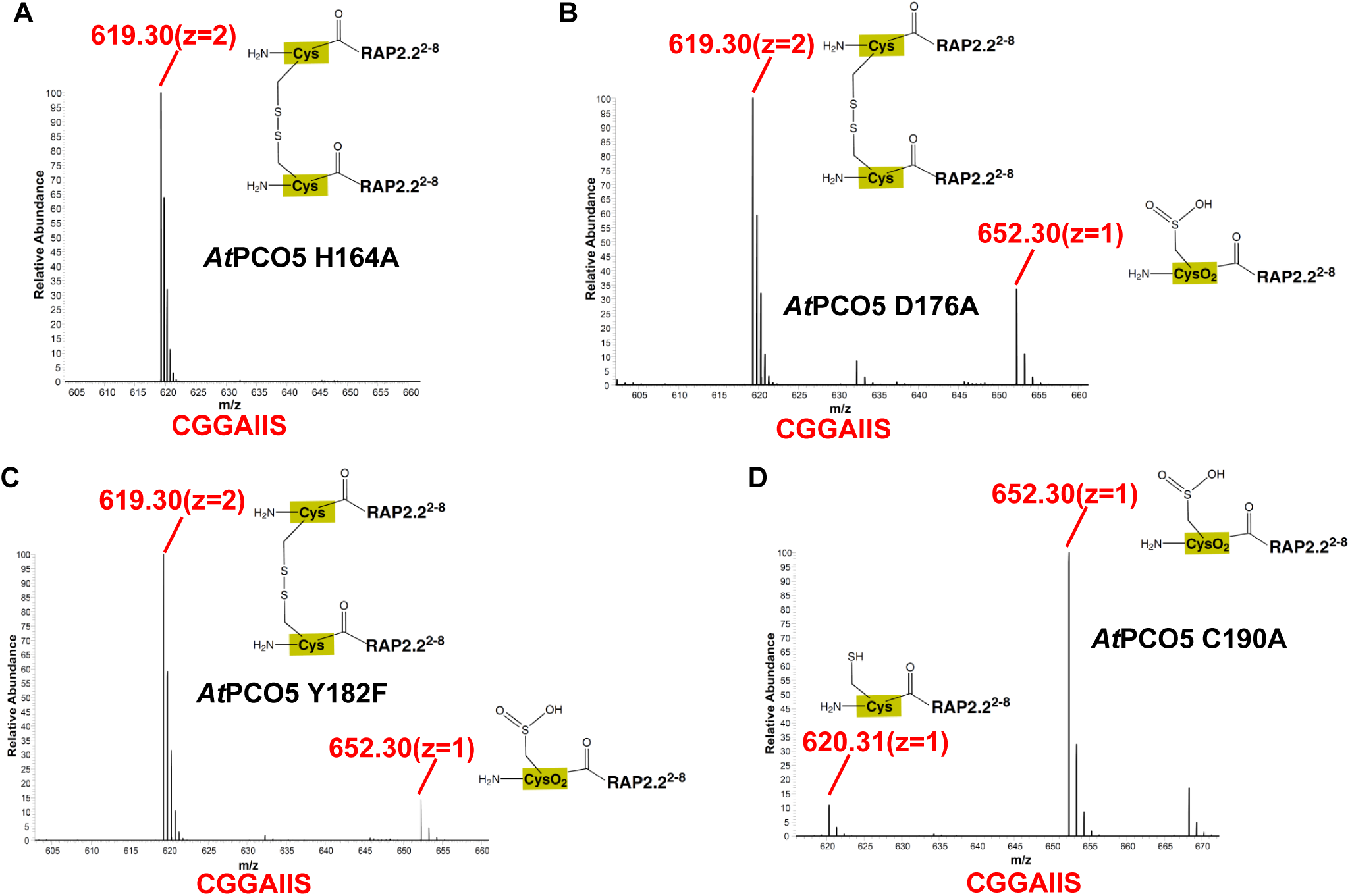
Mass spectra data of peptide AtRAP2.2^2-8^ (CGGAIIS) catalyzed by the variants of *At*PCO5. Spectra of peptide catalyzed by (A) *At*PCO5_1-242_ H164A, (B) *At*PCO5_1-242_D176A, (C) *At*PCO5_1-242_ Y182F, and (D) *At*PCO5_1-242_ C190A.

## Notes

### Competing Interest Statement

The authors have declared no competing interest.

